# Antibiotic killing of diversely generated populations of non-replicating bacteria

**DOI:** 10.1101/464628

**Authors:** Ingrid C. McCall, Nilang Shah, Adithi Govindan, Fernando Baquero, Bruce R. Levin

## Abstract

Non-replicating bacteria are known to be (or at least commonly thought to be) refractory to antibiotics to which they are genetically susceptible. Here, we explore the sensitivity to killing by bactericidal antibiotics of three classes of non-replicating populations of planktonic bacteria; (1) stationary phase, when the concentration of resources and/or nutrients are too low to allow for population growth; (2) persisters, minority subpopulations of susceptible bacteria surviving exposure to bactericidal antibiotics; (3) antibiotic-static cells, bacteria exposed to antibiotics that prevent their replication but kill them slowly if at all, the so-called bacteriostatic drugs. Using experimental populations of *Staphylococcus aureus* Newman and *Escherichia coli* K12 (MG1655) and respectively 9 and 7 different bactericidal antibiotics, we estimate the rates at which these drugs kill these different types of non-replicating bacteria. Contrary to the common belief that bacteria that are non-replicating are refractory to antibiotic-mediated killing, all three types of non-replicating populations of these Gram-positive and Gram-negative bacteria are consistently killed by aminoglycosides and the peptide antibiotics, daptomycin and colistin, respectively. This result indicates that non-replicating cells, irrespectively of why they do not replicate, have an almost identical response to bactericidal antibiotics. We discuss the implications of these results to our understanding of the mechanisms of action of antibiotics and the possibility of adding a short-course of aminoglycosides or peptide antibiotics to conventional therapy of bacterial infections.

## Introduction

For therapeutic purposes, the relationship between the concentrations of antibiotics and the rates of growth and death of bacteria, pharmacodynamics, is almost exclusively studied *in-vitro* under conditions that are optimal for the action of these drugs; relatively low densities of bacteria growing exponentially in media and under culture conditions where all members of the exposed population have equal access to these drugs, resources, wastes and metabolites excreted into the environment. To be sure, in some sites and tissues in acutely infected hosts, relatively low densities of the target pathogens may be growing exponentially at their maximum rate and thus are under conditions that are optimal for the action of antibiotics. However, this situation is almost certainly uncommon in established, symptomatic and thereby treated infections where the offending bacteria are likely to be compartmentalized in different sites and tissues and confronting the host’s immune defenses (1).

Infecting populations of bacteria may be non-replicating for different reasons and by different mechanisms. First, they may have exhausted the locally available resources; thus modified their environment so their populations are at or near stationary phase (2–6). Second, although local nutrients may be sufficient for their replication, for hosts treated with bactericidal drugs these bacteria may be minority populations of physiologically refractory survivors, the so-called “persisters” (7–13). Third, the offending bacteria may be non-replicating because of exposure to bacteriostatic antibiotics, a state we shall refer to as antibiotic-induced stasis. Fourth, infecting bacteria may be slowly replicating or at stationary phase inside phagocytes or other host cells (14, 15), or attached to the surfaces of tissues or prosthetic devices and within polysaccharide matrices know as biofilms (16) and thereby not replicating for one or more of previously described reasons (4, 17).

The concept of antibiosis has been classically linked to the fight against of microbes invading host tissues and thereby implying their active replication. With some exceptions associated with permeability (18), the susceptibility of bacteria to killing by bactericidal antibiotics is related to their rate of replication. In fact, with beta-lactams, the rate at which bacteria are killed has been shown to be strictly proportional to the rate at which the population is growing (19, 20). The same trend seems to occur for other bactericidal agents as fluoroquinolones, aminoglycosides, glycopeptides, and lipopeptides (21–24). It is well known that exposure to bacteriostatic antibiotics markedly reduce the efficacy of beta-lactam drugs to kill bacteria (25–27). However, save for these cases of antagonism between bacteriostatic and bactericidal drugs and the now more than a quarter of century old classical studies by R. Eng and colleagues (28), despite the potential clinical implications, there is remarkable little information about the pharmacodynamics of antibiotics for non-replicating populations of bacteria.

In this investigation we address two fundamental questions about the pharmacodynamics of non-replicating bacteria. What antibiotics and to what extent do these drugs kill nonreplicating bacteria? With respect to their susceptibility to antibiotic-mediated killing, are bacteria in these different non-replicating states physiologically similar? To address these questions we compare the activity of antibiotics on non-replicating bacterial populations obtained by different procedures. We present the results of experiments estimating the susceptibility of various non-replicating populations of *S. aureus* and *E. coli* to killing by respectively 9 and 7 different bactericidal antibiotics. We consider three types of non-replicating states of planktonic bacteria; (i) those at stationary phase in oligotrophic culture, (ii) the non-replicating survivors of exposure to bactericidal antibiotics, persisters, and (iii) bacteria exposed to bacteriostatic antibiotics, antibiotic-static populations. Contrary to the popular conception that antibiotics are ineffective at killing bacteria that are not replicating (19, 29, 30), albeit not the beta-lactam drugs, even at relatively low concentrations a number of existing bactericidal antibiotics can kill non-replicating bacteria of all three states. The results of our experiments indicate that the same classes of antibiotics, the aminoglycosides and the peptides, are particularly effective at killing non-replicating bacteria irrespective of the mechanism responsible for their failure to replicate. In addition to being relevant clinically, these results are interesting mechanistically; they suggest non-replicating bacteria of different types may share a common cell physiology with respect to their interactions with antibiotics.

## Results

### Antibiotic-mediated killing of exponentially growing bacteria

As a baseline for our consideration of the antibiotic susceptibility of non-replicating bacteria, we explore the response of exponentially growing populations *S. aureus* and *E coli* MG1655 to antibiotics. The results of these experiments are presented in Figure 1.

**Figure 1.**
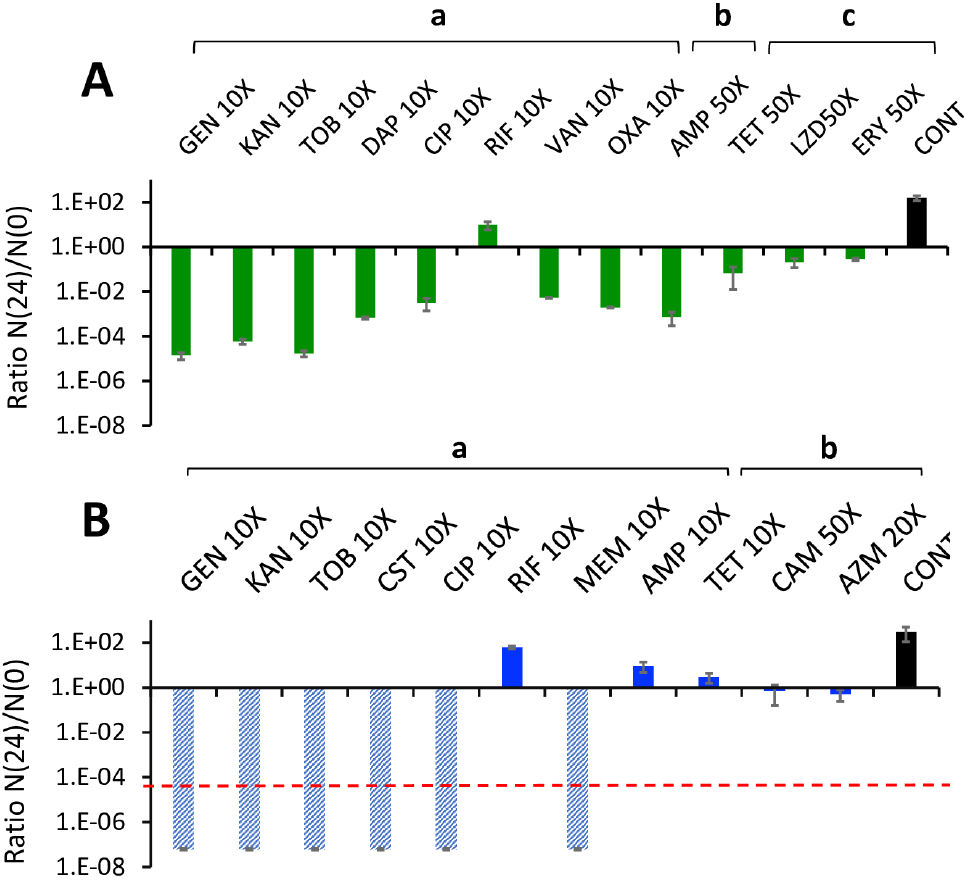
Antibiotic-mediated killing of exponentially growing cells. Ratio of the viable 24 hour and initial viable cell densities, N(24)/N(0), of exponentially growing broth cultures exposed to the concentrations of bactericidal and bacteriostatic antibiotics (multiples of MIC values). (**A**) *Staphylococcus aureus* Newman in MHII. (**B**) *Escherichia coli* MG1655 in LB. Mean and standard error of the N(24)/N(0) ratios from three independent experiments each with three samples. The broken line is the limit of detection (10^2^ cells per ml, hatched bars mean that this limit was surpassed in the assay). CONT is the control, the bacteria introduced into antibiotic free medium. The initial, N(0), densities of *S. aureus* and *E. coli* in these experiments are listed in the table below.

|  | a | b | c |
| --- | --- | --- | --- |
| *S. aureus* (A) | $1.25 \pm 0.05 \times 10^7$ | $2.27 \pm 0.15 \times 10^8$ | $2.80 \pm 0.50 \times 10^7$ |
| *E. coli* (B) | $1.62 \pm 0.17 \times 10^7$ | $8.27 \pm 3.99 \times 10^7$ | - |

For *S. aureus*, the aminoglycosides, gentamicin, kanamycin and tobramycin kill to the greatest extent, with reductions in viable cell density of more than 4 orders of magnitude. Daptomycin, ciprofloxacin, vancomycin, oxacillin and ampicillin are clearly bactericidal and reduce the viable cell density by 2-3 orders of magnitude. The increase in the N(24)/N(0) ratio for rifampin can be attributed to the ascent of rifampin-resistant mutants. Even at 50X MIC, tetracycline, linezolid, and erythromycin are effectively bacteriostatic. When exponentially growing cultures of *E. coli* are exposed to 10X MIC of gentamicin, kanamycin, tobramycin, colistin, ciprofloxacin and meropenem, the viable cell density is below that which can be detected by plating. As with *S. aureus*, the failure of rifampin to reduce the viable cell density can be attributed to the ascent of rifampin-resistant mutants. For *E. coli* exposed to ampicillin, our results suggest that the decay in the effective concentration of this drug, probably due to the chromosomal beta-lactamase effect at high initial densities can account for the failure of this bactericidal antibiotic to reduce the viable cell density of *E. coli*. As evidence for this, when cell-free extracts of the 24-hour ampicillin cultures are spotted onto lawns of sensitive *E. coli* MG1655, there is no zone of inhibition as there is when we spot the original media containing 10XMIC ampicillin.

### 2- Antibiotic-mediated killing of stationary phase bacteria

For the stationary phase experiments with both *S. aureus* and *E. coli*, we used cultures that had been incubated under optimal growth conditions for 48 hours. To estimate the amount of unconsumed, residual resources in these 48-hour stationary phase cultures, and thereby the capacity for additional growth, we centrifuged and filtered (0.20 microns) 48-hour cultures of these bacteria. We then added, ~10^3^ cells from overnight cultures to the cell-free filtrates, and estimated the initial viable cell density after 24 hours of incubation, respectively N(0) and N(24). The results of these experiments with four independent *E. coli* cultures and three independent *S. aureus* cultures, suggested no significant growth for *E. coli*, an N(24)/N(0) ratio of 0.36 ±0.22 and some growth for *S. aureus* an, N(24)/N(0) ratio of 10.91 ± 5.13. It should be noted however, that absence of and with limited growth in these spent media may be a reflection of the increase in pH in these media from pH7 at time 0 to pH 8.5 at 48 hours (31)

At 48 hours, the viable cell densities of the stationary phase cultures were estimated (As shown in Materials & Methods). In Figure 2, we present the results of this stationary phase experiment.

**Figure 2—.**
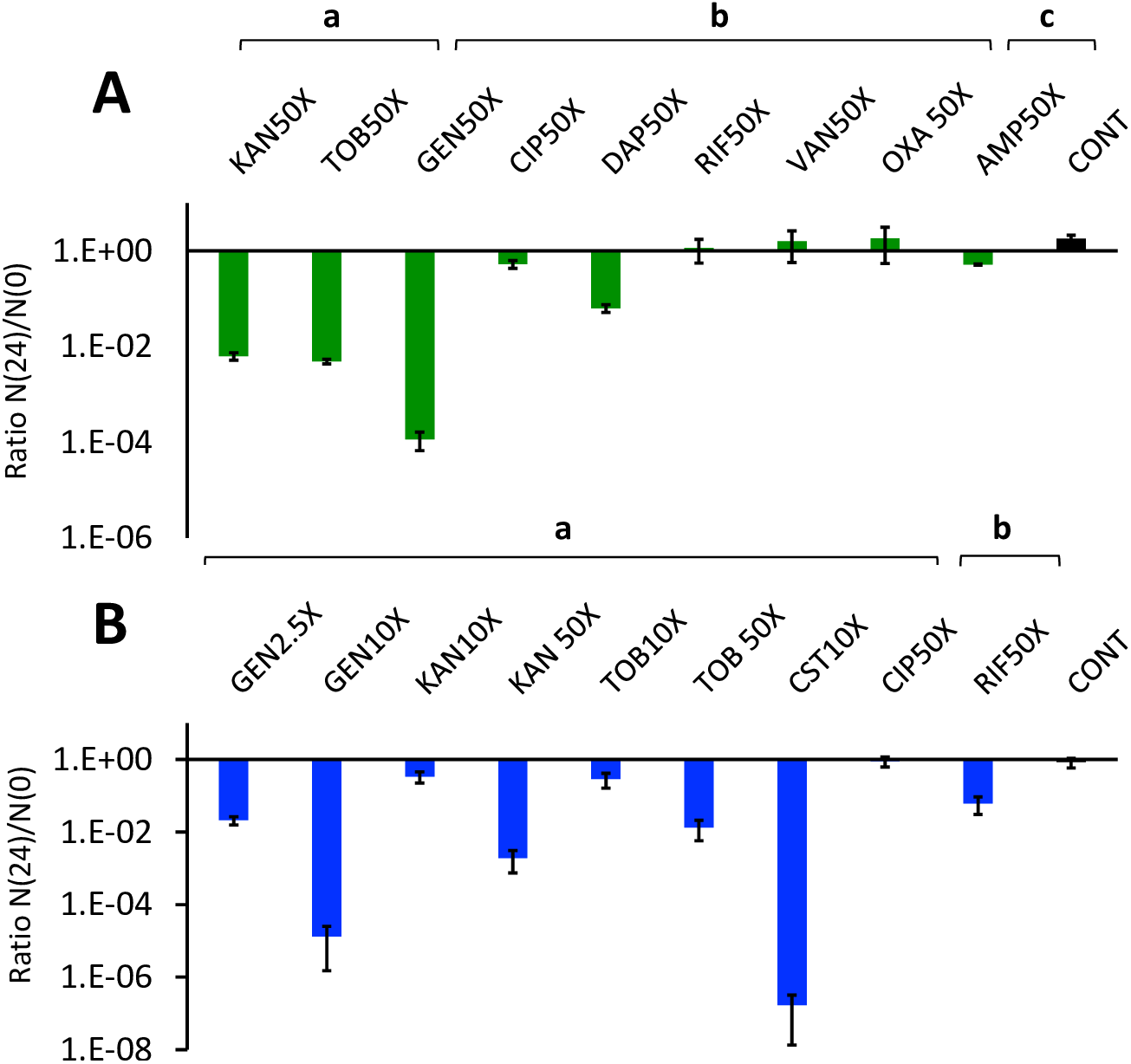
Antibiotic treatment of stationary phase bacteria. Ratio of the viable cell densities after and before twenty-four hours of exposure to bactericidal antibiotics, N(24)/N(0). Forty-eight-hour stationary phase cultures of *S. aureus* (**A**) and *E. coli* (**B**) were treated with the noted multiples of MIC of different drugs and antibiotic free controls (CONT). Mean and standard error of the N(24)/N(0) ratios from five and eleven independent experiments for (**A**) and (**B**), respectively. The mean and standard errors of the initial viable cell densities, N(0), in these experiments are listed in the table below.

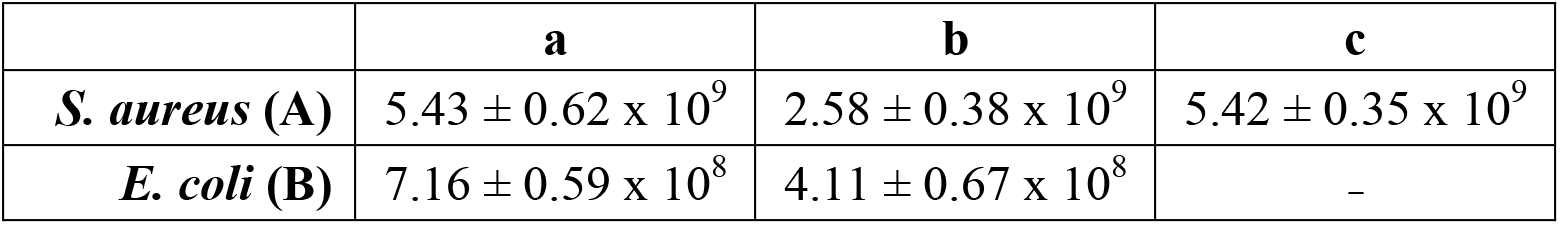

|  | a | b | c |
| --- | --- | --- | --- |
| *S. aureus* (A) | $5.43 \pm 0.62 \times 10^9$ | $2.58 \pm 0.38 \times 10^9$ | $5.42 \pm 0.35 \times 10^9$ |
| *E. coli* (B) | $7.16 \pm 0.59 \times 10^8$ | $4.11 \pm 0.67 \times 10^8$ | - |

In the absence of treatment (the control), there is no significant mortality between 48 and 72 hours for either *S. aureus* or *E. coli*. For *S. aureus*, only high concentrations of the aminoglycosides, gentamicin, kanamycin and tobramycin, and the cyclic peptide, daptomycin are effective in reducing the viable density of these 48-hours stationary phase cultures (Figure 2A). For *E. coli*, the aminoglycosides, gentamicin, tobramycin, and kanamycin are also effective for killing stationary phase cells, as is colistin. There is no evidence for the other bactericidal antibiotics tested, ciprofloxacin and rifampin killing stationary phase *E. coli*.

### 3) Antibiotic-mediated killing of *E. coli hip*A7 and *S. aureus* bactericidal persisters

Although density of exponentially growing *E. coli* surviving exposure to 10XMIC ampicillin in the experiment presented in Figure 1 is in excess of 10^9^, these *E. coli* are not persisters, but rather ampicillin sensitive cells that grew following the β-lactamase - mediated reduction in the effective concentration of ampicillin. In pilot experiments, where we increased the concentration of ampicillin to 25XMIC, the viable cell density of surviving *E. coli* was too low to test for the susceptibility of these bacteria to killing by other antibiotics. To address this issue, we restricted our *E. coli* persister experiments to *hipA7* (the Moyed mutant (32)) a construct that produces 10^3^ −10^4^ times greater numbers of persisters than wild type due to an increase in the basal level of (p)ppGpp synthesis (33). This is illustrated in Figure 3, where we compare the dynamics of formation and the relative densities of persisters for *E. coli* MG1655 and the *hipA7* construct exposed to 10X MIC ampicillin.

**Figure 3—.**
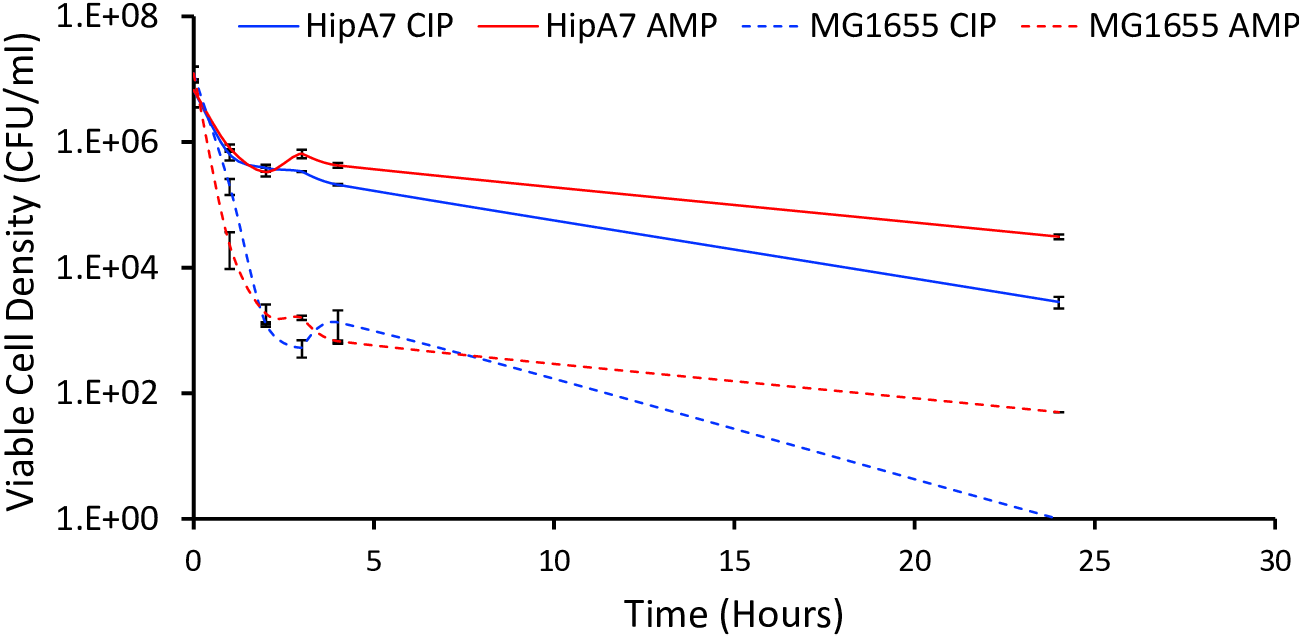
Dynamics of *hipA7* persister formation. Changes in viable cell density of exponentially growing *E. coli* MG1655 (broken lines) and a *hipA7* mutant of this strain (solid lines) exposed to 10X MIC ampicillin (red) and 10X MIC ciprofloxacin (blue). All data points represent the mean and standard error of three independent measurements. Average initial cell densities were 1.23 ± 0.34 x 10^7^ CFU/ml and 6.71 ± 3.16 x 10^6^ CFU/ml for MG1655 and *hipA7*, respectively.

For *S. aureus*, we restricted our experiments exploring the sensitivity of persisters to treatment with bactericidal antibiotics to ampicillin-generated persisters. In our pilot experiments generating these persisters with ciprofloxacin, kanamycin and tobramycin the viable cell density of surviving cells was too low to test for the susceptibility to killing by other antibiotics. To generate these *S. aureus* persisters, we used a protocol similar to that employed by (34). The results of these experiments are presented in Figure 4A. The extent to which these ampicillin-exposed *S. aureus* die following subsequent treatment is noted in by the N(24)/N(0) ratios of the controls, CONT. These *S. aureus* persisters are refractory to killing at 50XMIC concentrations of the bactericidal antibiotics ciprofloxacin, vancomycin, oxacillin and 20XMIC rifampin. This is not the case for the aminoglycosides, 5XMIC gentamicin, and 20XMIC tobramycin and kanamycin reduce the viable cell densities of these persisters by nearly three orders of magnitude. Albeit to an extent less than these aminoglycosides, at 20XMIC, the cyclic peptide daptomycin also kills these ampicillin-generated S. *aureus* persisters.

**Figure 4—.**
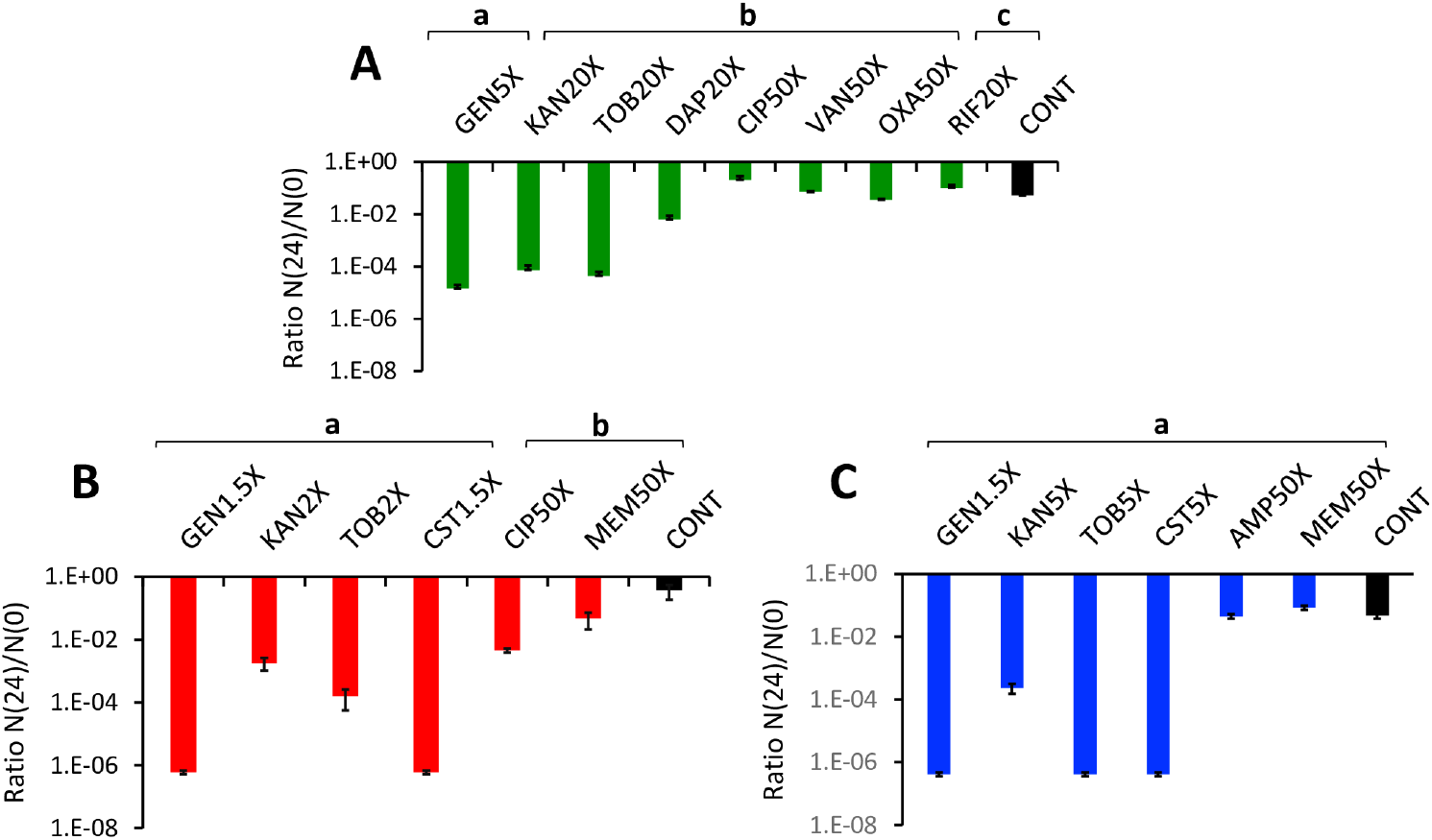
Antibiotic-mediated killing of bactericidal persisters. Ratio of viable cell density of bactericidal persisters after and before twenty-four hours exposure to the noted concentrations of bactericidal antibiotic, N(24)/N(0). (**A**) *S. aureus* treated with 25X MIC ampicillin to generate the persisters N(0). (**B**) *E. coli hipA7* treated with 10X MIC ampicillin N(0). (**C**) *E. coli hipA7* treated with 10X MIC ciprofloxacin to generate the persisters N(0). Mean of and standard error of the N(24)/N(0) ratios of three independent experiments. The mean and standard error of the N(0) densities of these experiments are presented in the below table..

|  | a | b | c |
| --- | --- | --- | --- |
| *S. aureus* (A) | $4.08 \pm 0.07 \times 10^7$ | $2.04 \pm 0.36 \times 10^7$ | $1.14 \pm 0.09 \times 10^7$ |
| *E. coli* (B) | $1.72 \pm 0.24 \times 10^6$ | $4.37 \pm 2.61 \times 10^6$ | - |
| *E. coli* (C) | $5.06 \pm 0.84 \times 10^6$ | - | - |

The *hipA7* persisters were prepared by exposure to ampicillin and ciprofloxacin with a protocol similar to that employed by Keren and colleagues(35). The results of these experiments are presented in Figures 4B and 4C. Relative to the controls, low, but super-MIC concentrations of the aminoglycosides, gentamicin, kanamycin and tobramycin and also the peptide colistin reduce the viable cell density of the *E. coli hipA7* persisters by three to six orders of magnitude. Even at 50XMIC the other antibiotics have little or no effect in reducing the viable cell density of the *E. coli hipA7* persisters.

### 4- Antibiotic-mediated killing of antibiotic-induced static populations

Antibiotic-induced static populations were generated by exposing exponentially growing *S. aureus* and *E. coli* to bacteriostatic drugs for 24 hours, followed by second antibiotics for another 24 hours (as shown in Materials & Methods). The results of these experiments are presented in Figure 5.

**Figure 5—.**
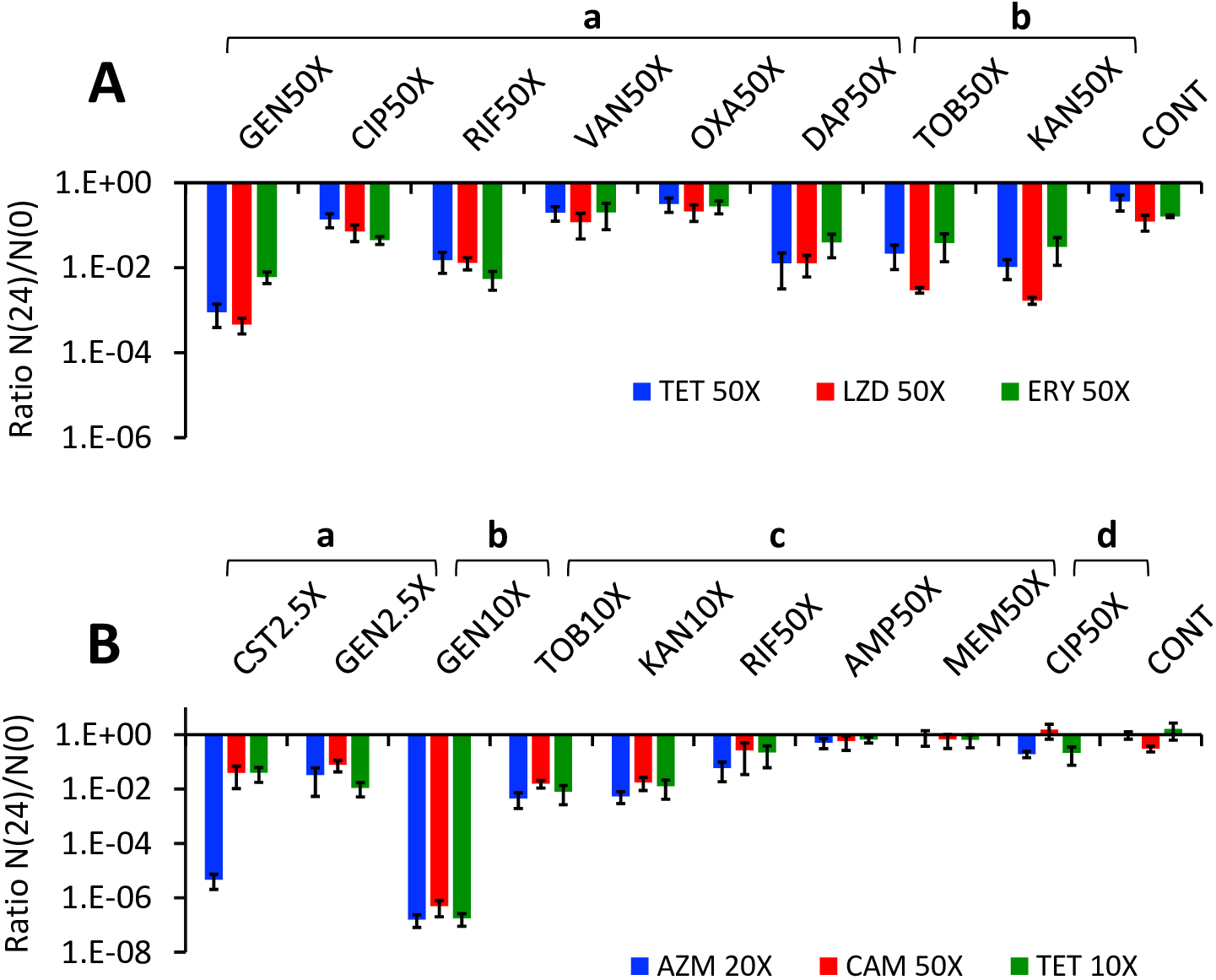
Antibiotic-mediated killing of antibiotic-static populations. Mean and standard error of the N(24)/N(0) ratios (**A**) *S. aureus*, (**B**) *E. coli*. The concentrations of the antibiotics as multiples of MICs used to generate the static populations and for subsequent treatment are noted in the figure. The initial densities of the *S. aureus* antibiotic-static populations for tetracycline (TET), linezolid (LZD), erythromycin (ERY) are shown below in the *S. aureus* table. For the *E. coli* antibiotic-static populations initial densities for azithromycin (AZM), chloramphenicol (CAM) and tetracycline (TET) are shown in the *E. coli* table below.

| *S. aureus* | a | b | c | d |
| --- | --- | --- | --- | --- |
| **TET** | $4.24 \pm 1.28 \times 10^7$ | $1.04 \pm 0.22 \times 10^7$ | - | - |
| **LZD** | $4.16 \pm 1.60 \times 10^7$ | $3.83 \pm 0.57 \times 10^6$ | - | - |
| **ERY** | $2.71 \pm 1.13 \times 10^7$ | $1.23 \pm 0.26 \times 10^7$ | - | - |

| *E. coli* | a | b | c | d |
| --- | --- | --- | --- | --- |
| **AZM** | $1.28 \pm 0.45 \times 10^7$ | $1.85 \pm 0.69 \times 10^7$ | $1.74 \pm 0.43 \times 10^7$ | $1.13 \pm 0.41 \times 10^7$ |
| **CAM** | $6.56 \pm 1.57 \times 10^6$ | $7.56 \pm 2.96 \times 10^6$ | $6.39 \pm 1.48 \times 10^6$ | $4.88 \pm 1.15 \times 10^7$ |
| **TET** | $2.72 \pm 0.34 \times 10^7$ | $3.76 \pm 0.56 \times 10^7$ | $2.44 \pm 0.68 \times 10^7$ | $6.55 \pm 1.24 \times 10^7$ |

The *S. aureus* antibiotic-static populations surviving exposure to tetracycline, erythromycin and linezolid are killed by the aminoglycosides, rifampin and daptomycin. The *E. coli* antibiotic-static populations surviving exposure to azithromycin, chloramphenicol and tetracycline are readily killed by the aminoglycosides and colistin and marginally if at all so by high concentrations of rifampin. Higher concentrations of gentamicin (10XMIC) are more effective than 10XMIC of the other aminoglycosides.

## Discussion

During the course of an infection, several different conditions contribute to the nonreplicating status of initially growing populations of the infecting bacteria, triggering type I persisters, those slow growers that are generated by exposure to a stress signal (36). Among these conditions are stationary phase resulting from a dearth of the nutrients and resources in the infected tissues, the formation of biofilms, and the immune defenses, primarily engulfment by professional and amateur phagocytic host cells. Therapy with antibiotics will also result in the production of non-replicating populations of bacteria; persisters surviving exposure to bactericidal antibiotics, and stasis induced by bacteriostatic drugs. We postulate that although they are generated in different ways, common mechanisms are responsible for the failure of these bacteria to replicate and thereby for the same classes of antibiotic to kill them. These mechanisms involve the general stringent response, a strongly regulated process governed by the alternative sigma factor RpoS (35) up-regulated by the accumulation of hyperphosphorylated guanine nucleotides, as the “alarmone” (p)ppGpp (37–39). That regulation might in fact reflect the metabolism of persisters which is oriented toward energy production, with depleted metabolite fluxes, which perpetuates the slow growth (12, 40).

The contribution of the stringent response to antibiotic-induced stress is not well understood. A number of observations suggest that antibiotic exposure can trigger a RpoS-stationary response and the generation of non-replicating populations. Sub-inhibitory concentrations of beta-lactams induce the stringent response (41, 42). Response to stress by bacteriostatic antibiotics acting on the ribosome (as macrolides, chloramphenicol, or tetracyclines) is likely to be manifest by the reduction in protein synthesis, which might reduce the building-up of (p)ppGpp, but that might be overcompensated by reduced degradation of this nucleotide (43, 44). Proteome analysis of erythromycin-exposed “permeable” *E. coli* suggests a RpoS-regulated profile (45). In fact, sub-inhibitory concentrations of bacteriostatic antibiotics induce the stringent response, leading to beta-lactam tolerance (46).

Cationic peptides, including polymyxins, do not elicit RpoS response, but rather increase the permeability of the cell membranes and thereby act as rapid “external killers” (47, 48). Whether the aminoglycosides can elicit an RpoS response is unclear at this time. As is the case of other protein synthesis inhibitors, this machinery might be less effective in the presence of the antibiotic, than in the absence; the same occur with oxidative stress responses (44). The rapid killing observed in our work (see Figure 1) may well be because the stringent response has no time to develop when exposed to these drugs. However, if aminoglycosides are able to induce the RpoS response, that might favor aminoglycoside transport into cells that results in membrane-damage and killing of nongrowing cells (49).

The cellular immune response might also be responsible for generating non-replicating populations. Salmonella RpoS-dependent genes are activated into the intracellular environment of eukaryotic cells (50). An RpoS activated system contributes to survival of *E. coli* and *Salmonella* to the phagocyte oxidase-mediated stress resistance (51, 52) and influences intracellular survival of *Legionella* in macrophages and in *Acanthamoeba* cells (53–55). Intracellular survival is also related with the overexpression of heat-shock stress proteins, promoting non-replication (56, 57).

These studies suggest that non-replicating bacteria might also arise by the stringent response derived from professional and/or non-professional phagocytosis. Of course, aminoglycosides and peptide antibiotics do not enter in eukaryotic cells, but nonreplicating bacteria can be released from phagocytes lysed by bacteria-induced programmed necrosis, contributing to the chronification of the infection (58). Antibody-antibiotic conjugates might improve therapy of phagocytized bacteria where aminoglycosides and peptides are excluded (15).

We postulate that essentially the same set of cellular responses occurs in the different stress-inducing conditions that bacteria encounter during the infection process, and different classes of antibiotics are similarly effective in killing these different types of non-replicating cells. These effects probably are expressed as an hormetic (dose-dependent) stress response(59, 60). Of course, non-growth is a complex mechanism in which the final antibiotic effects are influenced by those resulting from different coexisting stresses, for instance, non-growth resulting from ribosome hibernation could facilitate some degree of gentamicin tolerance (61). The non-growing phenotype is therefore causally complex, which negatively influences the full quantitative repeatability of the experiments.

It is well known that stationary phase populations of bacteria are refractory to killing by some antibiotics (19, 29, 30). The results presented here illustrate that this is not the case for other antibiotics, and particularly some aminoglycosides and peptide antimicrobials kill stationary phase populations of *E. coli* and *S. aureus*. This observation is not new; it has been known for some time that the aminoglycosides and the cyclic peptides, daptomycin and colistin are capable of killing stationary phase bacteria (24, 61–63). Less seems to be known about the susceptibility to killing by bactericidal antibiotics of the other non-replicating states of bacteria considered here, persisters and bacteria with antibiotic-induced stasis.

Although there have been many studies of persistence, relatively little is known about the susceptibility of these non-replicating bacteria to antibiotics other than those employed to generate them. One exception to this is a study by Allison and colleagues, (64) that demonstrates that by adding metabolites, *E. coli* and *S. aureus* persisters in the form of biofilms become susceptible to killing by aminoglycosides. Our results with the ampicillin-generated planktonic *S. aureus* persisters are fully consistent with these observations. Results with the *hipA7* persisters used in this work also suggest that *E. coli* “natural” persisters are sensitive to killing by even low concentrations of the aminoglycosides and the peptide colistin.

There is also a relative dearth of quantitative information about the susceptibility to antibiotic-mediated killing of non-replicating bacteria induced by exposure to bacteriostatic antibiotics. To be sure, within the first decade after the discovery of antibiotic, there was evidence for exposure to bacteriostatic drugs reducing the efficacy of bactericidal (65) and these observations were corroborated more recently (27). Early observations concerning the lower efficacy of penicillin in static cells produced by chloramphenicol and the tetracyclines have engendered what some may see as an immutable law in the practice of antibiotic therapy, do not mix bacteriostatic and bactericidal drugs. However, we show that some existing bactericidal antibiotics are quite effective in killing bacteria that are not replicating because exposure to bacteriostatic antibiotics, what we refer to as antibiotic-static populations.

In summary, stationary-phase *S. aureus* populations are killed at a substantial rate by the aminoglycosides and to a lesser extent by daptomycin. *S. aureus* persisters generated by exposure to ampicillin are also killed aminoglycosides and the lipopeptide daptomycin, but not the other drugs tested. Antibiotic-induced static populations of *S. aureus* maintain the same killing profile, with aminoglycosides and daptomycin as the sole killing agents, with the exception, in this case of rifampicin. Some aminoglycosides and the peptide colistin are also effective at killing stationary phase *E. coli*, whilst the other antibiotics tested were not. This is also the case for the *hipA7 E. coli* persister and *E. coli* “suffering” from the stasis induced by ribosome-targeting bacteriostatic antibiotics. This homogeneous pattern of response to antibiotic killing in differently originated slow-growth populations support a basically homogeneous physiology in all type I persisters.

### Potential clinical implications

In recent years there has been a great deal of interest in discovering and developing drugs to treat non-replicating populations of bacteria, particularly those associated with biofilms. A prime example of this is Kim Lewis and colleague’s (66) use of a novel antibiotic, acyldepsipeptide (ADEP4). Despite growing efforts in the endeavor to find new antibiotics to kill non-replicating bacteria, the results presented here suggest that existing antimicrobials may well be up to that task. A well-warranted concern is that the antibiotics with this virtue are among the more toxic drugs, aminoglycosides and the peptides (67–69). It should be noted, however, relatively low, and possible non-toxic concentrations of the aminoglycosides and the peptide, colistin can kill *E. coli* antibiotic-static and *hipA7* persisters. Most importantly, as has been the case with cancer chemotherapy, there are conditions under which some toxic side-effects of systemic treatment are more then made up for by the sometimes life-saving benefits of that treatment (70). Of course, inhaled therapy, providing very high local concentrations of aminoglycosides or peptidic antibiotics, has proven its efficacy in mostly non-growing populations of *Pseudomonas aeruginosa* and *Staphylococcus aureus* involved in chronic lung colonization in cystic fibrosis patients (71). Also, high local concentrations of these antibiotics have been useful in intravesical therapy of recurrent urinary tract infections (72) or in catheter locking solutions to treat catheter-related bloodstream infections (73).

How important persisters are clinically is, at this juncture, not all that clear. Persisters remain susceptible to phagocytosis and other elements of the innate immune system, the main factor influencing control of infections, can be attributed to the innate immune system, and they would play little or no role in reducing the efficacy of antibiotic therapy (74). Consistent with this yet-to-be tested (but testable) hypothesis in experimental animals and patients is the observation that for immune competent patients, bacteriostatic drugs are as effective for treatment as highly bactericidal agents (75, 76).

As intriguing as they may be scientifically, planktonic persisters surviving exposure to bactericidal antibiotics are not the majority of the non-replicating bacteria present during the infection process. Stationary phase bacteria resulting from local shortage of nutrients, non-growing populations induced by bacteriostatic agents, biofilm populations and phagocytosed bacteria (eventually released by the lysis of the engulfing phagocytes) are likely to be the majority of phenotypically antibiotic resistant bacteria in an established infection. They are certainly involved in chronification of infections, and subsequent reactivations and relapses. It may well be that along with the standard therapy, the addition of a short-course administration of antibiotics that kill these nonreplicating bacteria, like the aminoglycosides and peptides, may well accelerate the course of treatment and increase the likelihood of its success.

## Materials & Methods

### Bacteria

*Staphylococcus aureus* Newman obtained from Dr. William Shafer, Emory University, *E. coli* K12 MG1655 obtained from Ole Skovgaard, Roskilde University, and the high frequency persister strain of E. coli K12, *hipA7* constructed by Moyed and Betrand (32) obtained from Kyle Allison, Emory University.

### Liquid culture

For the *S. aureus* Mueller-Hinton Broth (MHII) (275710, BD™) and for *E. coli* Lysogeny Broth (LB) (244620, Difco). These antibiotic-kill experiments were performed in 6 well polystyrene plates, CELLTREAT™.

### Sampling Bacterial Densities

The densities of bacteria were estimated by serial dilution in 0.85% saline and plating on LB (1.6%) agar plates.

### Antibiotics and their sources

Ampicillin, chloramphenicol, colistin, gentamicin, kanamycin, oxacillin, tetracycline, and vancomycin – SIGMA, azithromycin and tobramycin, TOCRIS, daptomycin, TCI, erythromycin, MP BIOCHEMICALS, ciprofloxacin and rifampin, APPLICHEM, meropenem-COMBI-BLOCKS, linezolid – CHEM IMPEX INT’L

### Minimum Inhibitory Concentrations (MICs)

*For* both *S. aureus* Newman and *E. coli* MG1655 the MICs of the antibiotic were estimated using the two-fold micro dilution procedure (77). Two different initial concentrations of antibiotics were used to obtain accurate measurements from the two-fold micro dilution procedure. The estimated MICs of each of the antibiotic bacteria combination are listed in Table 1.

**Table 1.** Minimum Inhibitory Concentrations in μg/ml.

| Antibiotic | *S. aureus* Newman | *E. coli* MG1655 | *E. coli* MG1655<br>*hipA7* |
| --- | --- | --- | --- |
| Ampicillin (AMP) | 4.7 | 4.7 | 4.7 |
| Azithromycin (AZM) | 2.3 | 19.0 | 38.0 |
| Chloramphenicol (CAM) | 13 | 4.7 | 4.7 |
| Ciprofloxacin (CIP) | 0.2 | 0.6 | 0.6 |
| Colistin (CST) | N/A | 1.1 | 1.1 |
| Daptomycin (DAP) | 1.6 | N/A | N/A |
| Erythromycin (ERY) | 0.6 | N/A | N/A |
| Gentamicin (GEN) | 0.8 | 9.3 | 9.3 |
| Kanamycin (KAN) | 2.3 | 4.7 | 4.7 |
| Linezolid (LZD) | 1.1 | N/A | N/A |
| Meropenem (MEM) | N/A | 1.2 | **0.6** |
| Oxacillin (OXA) | 0.3 | N/A | N/A |
| Rifampin (RIF) | 0.002 | 9.4 | 9.4 |
| Tetracycline (TET) | 0.6 | 2.3 | 75.0 |
| Tobramycin (TOB) | 0.3 | 1.2 | 2.3 |
| Vancomycin (VAN) | 1.6 | N/A | N/A |

### N(24)/N(0) Ratios

As our measure of the efficacy of the different antibiotics to killing the exposed bacteria, we use the ratio of the viable cell density after 24 hours of exposure to the drug and the initial density estimated prior to exposure, N(24)/N(0). For each experiment, we estimated the N(0) and N(24) densities (CFUs) with three independent serial dilutions. For each antibiotic - bacteria combination, unless otherwise stated, we ran at least three independent experiments and calculated the mean and standard error of the N(24)/N(0) ratios.

### Antibiotic-mediated killing of exponentially growing bacteria

For these experiments, overnight cultures of *S. aureus* and *E. coli* MG1655 were added to broth at a ratio of 1:100 and incubated for 1.5 hours and the density of the cultures estimated, N(0). Five mls of these cultures were then put into 6-well plates and the antibiotics added. The cultures with the antibiotics were incubated for 24 hours at which time the viable cell densities were estimated, N(24).

### Antibiotics-mediated killing of stationary phase bacteria

For the stationary phase experiments with both *S. aureus* and *E. coli*, we used cultures that had been incubated under optimal growth conditions for 48 hours. At 48 hours, the viable cell densities of the stationary phase cultures were estimated, N(0). Following that, five ml aliquots were put into 6-well plates, the antibiotics added, and the cultures incubated with shaking for another 24 hours, at which time the viable cell densities were estimated.

### Antibiotic-mediated killing of *S. aureus* bactericidal persisters

Overnight MHII cultures of *S. aureus* Newman were diluted 1/10 in fresh MHII and 25XMIC ampicillin added immediately. After 24 hours, the viable cell densities were estimated, N(0). Five mls of these ampicillin treated cultures were put into 6-well plates and the second antibiotic(s) added and the cultures incubated with shaking for another 24 hours.

### Antibiotic-mediated killing of *E. coli* hipA7 bactericidal persisters

Exponentially growing LB cultures of *E. coli hipA7* were exposed to 10X MIC ciprofloxacin or ampicillin for 4 hours, N(0) at which time they were treated with the second antibiotic for another 24 hours, N(24).

### Antibiotic-mediated killing of antibiotic-induced static populations

Antibiotic-induced static populations were generated by exposing exponentially growing *S. aureus* and *E. coli* to bacteriostatic drugs for 24 hours, at which time the viable cell density was estimated N(0). The culture was divided into five ml aliquots in 6-well plates and the second antibiotic added. The cultures were maintained for another 24 hours and the viable cell densities estimated, N(24).

#### A Caveat

As quantitative biologists, we found our experiments with antibiotic-mediated killing and persisters to be somewhat frustrating. The magnitude of the variation in the extent of antibiotic-mediated killing and levels of persistence among independent replicas was often substantial. We did a great deal of replication and are confident about the results reported here, in a semi-quantitative way. By the latter, we mean that we are convinced that the relative extents of kill and levels of persistence by the antibiotics used here would be obtained in parallel experiments in other laboratories. On the other hand, we would be surprised if the absolute numbers of bacteria surviving exposure to these drugs in these experiments would be identical to those reported here.

Curiously, when we discuss experimental studies of persistence and related topics with colleagues working on these subjects, they too whine about the variation among experiments, but this variation is rarely reported, for an exception see (78). Contributing to this variation is, of course, differences among batches of media, and particularly broths. Moreover and perhaps more important is variation in the strains used by different laboratories and possibly even by the same laboratory at different times. Although designated, *E. coli* MG1655 or *S. aureus* Newman, every time those bacteria are re-cloned, there is a possibility of the random fixation mutations, which in the course of time can affect the phenotype of the cell line including declines in fitness, a phenomenon called Muller’s Ratchet (79). This accumulation of mutations can be seen as DNA sequence data as well as phenotypic variation among isolates of strains with the same name, see for example (80, 81). In addition, it seems clear that persistence and the response of bacteria to antibiotics are multi-causal phenotypes; as is the rule in complex systems, the quantitative reproducibility of experiments is frequently impaired, as small and difficult-to-control variations in the initial variables can quantitatively influence the final result (82)

## Acknowledgement

This research was funded from by a grant from the US National Institutes of Health, GM098175 to BRL, and Regional Government of Madrid (InGEMICS-C, S2017/BMD-3691, co-financed by the European Development Regional Fund [ERDF], “A Way to Achieve Europe”) to FB. We wish to thank Melony Ivey and Esther Lee for superb technical help.

The funders of this endeavor had no role in designing this study, data collection and interpretation, or the decision to submit this work for publication.

